# Anti-pausing activity of region 4 of the RNA polymerase σ subunit and its regulation by σ-remodeling factors

**DOI:** 10.1101/2020.08.11.244855

**Authors:** Konstantin Brodolin, Zakia Morichaud

## Abstract

The basal transcription factors of cellular RNA polymerases (RNAPs) stimulate the initial RNA synthesis via poorly understood mechanisms. Here, we explored the mechanism employed by the bacterial factor σ in promoter-independent initial transcription. We found that the RNAP holoenzyme lacking the promoter-binding domain σ4 is ineffective in *de novo* transcription initiation and displays high propensity to pausing upon extension of RNAs 3 to 7 nucleotides in length. The σ4 domain stabilizes short RNA:DNA hybrids and suppresses pausing by stimulating RNAP active-center translocation. The anti-pausing activity of σ4 is modulated by its interaction with the β subunit flap domain and by the σ remodeling factors AsiA and RbpA. Our results suggest that the presence of σ4 within the RNA exit channel compensates for the intrinsic instability of short RNA:DNA hybrids by increasing RNAP processivity, thus favoring productive transcription initiation. This “RNAP boosting” activity of the initiation factor is shaped by the the thermodynamics of RNA:DNA interactions and thus, should be relevant for any factor-dependent RNAP.

## INTRODUCTION

Transcription initiation by DNA-dependent RNA polymerase (RNAP) is the first and highly regulated step in gene expression (Ruff *et al*, 2015)(Saecker *et al*, 2011). After initiation of *de novo* RNA synthesis at promoters (initial transcription), RNAP fluctuates between promoter escape, which leads to productive RNA synthesis, and stalling at promoters, which leads to “abortive”, reiterative RNA synthesis, a general phenomenon observed for all RNAPs (Carpousis & Gralla, 1980)(Kubori & Shimamoto, 1996)(Durniak *et al*, 2008) (Murakami *et al*, 2002). Nascent RNA chains longer than 8 nucleotides (nt) form stable, 9-to 10-base pair (bp)-long RNA:DNA hybrids that are considered a hallmark of the productive elongation complex (Nudler *et al*, 1997)(Sidorenkov *et al*, 1998)(Kireeva *et al*, 2000) (Kostrewa *et al*, 2009)(Vassylyev *et al*, 2007a). Nascent RNA chains shorter than 9 nt form unstable hybrids with ssDNA templates and tend to dissociate from the RNAP active site (Metzger *et al*, 1993)(Carpousis & Gralla, 1980)(Sidorenkov *et al*, 1998). However, short RNAs (5-6 nt in length) can be stably bound in the paused initially transcribing complex (ITC) formed at the *lac*UV5 promoter (Brodolin *et al*, 2004)(Duchi *et al*, 2016)(Dulin *et al*, 2018). The initial transcription pause occurring after the synthesis of 6-nt RNA functions as a checkpoint on the branched pathway between productive and non-productive transcription (Dulin *et al*, 2018).

During RNA synthesis, RNAP performs a stepwise extension of the RNA chain by one nucleotide that is called nucleotide addition cycle (NAC) (reviewed by (Belogurov & Artsimovitch, 2019). During the NAC, the fist initiating nucleoside triphosphate (NTP) or the 3’ end of RNA occupies the product site (i-site) (pre-translocated state), while the incoming NTP enters in the substrate site (i+1-site). After formation of the phosphodiester bond, the RNA 3’ end moves from the i+1-site to the i-site (post-translocated state). The concerted translocation of RNA and DNA to the active site is controlled by the β’ subunit trigger loop that folds into the trigger helix upon transition from the pre-translocated to the post-translocated state (Toulokhonov *et al*, 2007b)(Zhang *et al*, 2010).

The basal transcription initiation factors (i.e. bacterial σ subunit, archaebacterial TFB, and eukaryotic TFIIB) stimulate the initial steps of RNA synthesis (Kulbachinskiy & Mustaev, 2006)(Werner & Weinzierl, 2005)(Bushnell *et al*, 2004)(Campbell *et al*, 2002)(Zenkin & Severinov, 2004)(Sainsbury *et al*, 2013). Specifically, the σ subunit region 3.2 hairpin loop (σ3.2-finger) contacts the template ssDNA strand at positions −4/-5 and controls its positioning in the active site (Pupov et al, 2010)(Tupin *et al*, 2010). The σ3.2-finger can indirectly modulate the priming of *de novo* RNA synthesis at promoters (Kulbachinskiy & Mustaev, 2006)(Pupov *et al*, 2014) and promoter-like DNA templates, such as the M13 phage origin (Zenkin & Severinov, 2004). The σ subunit may also exert an inhibitory effect on initial transcription. Indeed, the unstructured linker connecting domains σ3 and σ4 (formed by the σ regions 3.2 and 4.1) is located in the RNA-exit channel and represents a barrier for growing RNA chains. This linker is displaced by RNA upon promoter escape (Li *et al*, 2020). A clash between the σ3.2-finger and >4-nt RNA chains hinders RNA extension and may cause the formation of abortive RNAs, thus contributing to pausing during initial transcription (Murakami *et al*, 2002)(Basu *et al*, 2014)(Duchi *et al*, 2016)(Zhang *et al*, 2012)(Zuo & Steitz, 2015)

Binding of the regions σ3.2 and σ4.1 within the RNA exit channel takes place during assembly of the RNAP holoenzyme when the σ subunit undergoes the transition from the “closed” to the “open” conformation (Callaci *et al*, 1999). Recent single-molecule fluorescence resonance energy transfer studies demonstrated that in *Mycobacterium tuberculosis*, this transition is regulated by the activator protein RbpA (Vishwakarma *et al*, 2018) that interacts with the σ2 and σ3.2 domains (Boyaci *et al*, 2018). Whether RbpA can influence initial transcription has never been explored.

Several mechanisms to explain the σ3.2-finger stimulatory activity during initial transcription have been proposed: (1) stabilization of the template ssDNA in the RNAP active site (Zhang *et al*, 2012); (2) decreased K_m_ for 3’-initiating NTP binding in the substrate i+1-site (Kulbachinskiy & Mustaev, 2006); and (3) stabilization of short RNAs in the active site (Campbell *et al*, 2002)(Zenkin & Severinov, 2004)(Zenkin *et al*, 2006). The last mechanism was also suggested for the B-reader domain of TFIIB, which is the structural homologue of the σ3.2-finger (Bushnell *et al*, 2004)(Chen & Hampsey, 2004). As the σ subunit occludes the RNA path and contacts all principal regulatory domains of RNAP (β’-clamp, β-lobe, β-Flap), it may affect RNA synthesis in several ways: through ssDNA template positioning, RNA binding, or direct modulation of the RNAP domain motions.

Here, to discriminate among these different scenarios, we investigated how the σ subunit and RNA:DNA hybrid length affect branching between productive and non-productive RNA synthesis during initial transcription by two RNAPs from phylogenetically distant bacterial lineages: *Escherichia coli* (*Eco*RNAP) and *M. tuberculosis* (*Mtb*RNAP). Compared with *Eco*RNAP, *Mtb*RNAP presents several structure-specific features, particularly the lack of *Eco*-specific TL-insertion and the presence of the ∼90 amino acid-long *Actinobacteria*-specific insertion in the β’ subunit (β’-SI) (Lane & Darst, 2010)(Lin *et al*, 2017). To analyze directly the effects of the σ subunit on the RNAP catalytic site activity, we used promoter-less DNA scaffold templates (Tupin *et al*, 2010) (Zenkin & Severinov, 2004). DNA scaffolds have been widely used in structural studies on ITCs of bacterial and eukaryotic RNAPs (Cheung *et al*, 2011) (Liu et al, 2011). The scaffold model allows bypassing the complexity of promoter-dependent initiation that is strongly influenced by the promoter configuration and by the interactions of σ with promoter elements. When complexed with RNAP, scaffold DNA templates harbor a “relaxed” conformation lacking the topological stress observed in the transcription bubble due to DNA scrunching during initial transcription at promoters. Moreover, as DNA scaffolds lack non-template strand ssDNA, transcription initiation should be less affected by interaction with the core recognition element (CRE) (Vvedenskaya *et al*, 2014). We found that the promoter-binding domain σ4 (i.e. the structural homologue of the eukaryotic TFIIB B-ribbon), located ∼60 Å away from the active site, strongly stimulates RNAP translocation and stabilizes short RNA:DNA hybrids in the RNAP active site. The combination of these activities provides the basis for the initiation-to-elongation transition regulation by the auxiliary transcriptional factors that binds to the σ subunit.

## RESULTS

### The σ^70^ subunit is required for initial transcription from the promoter-less scaffold DNA template

To explore the role of the σ subunit in initial transcription we used two types of minimal DNA scaffold templates (**Fig. 1A**): a <u>S</u>hort <u>D</u>uplex <u>T</u>emplate (SDT), which included the 9-bp downstream DNA (dwDNA) duplex, and a <u>L</u>ong <u>D</u>uplex <u>T</u>emplate (LDT), which comprised the 18 bp dwDNA duplex. The dwDNA duplex of the LDT scaffold forms additional contacts with the β’ subunit residues 202-247 that stabilize the RNAP-scaffold complex (Kulbachinskiy *et al*, 2002)(Vassylyev *et al*, 2007a). Previously, we demonstrated that extension of the 3-nt RNA primer on SDT DNA is strongly stimulated by the σ^70^ subunit (Tupin *et al*, 2010). Here, we found that the RNAP core from *E. coli* (*Eco*RNAP) was inactive in *de novo* transcription initiation at SDT and LDT templates performed in the presence of [α-^32^P]-UTP, CTP and GTP (**Fig. 1B**). Conversely, the σ^70^-*Eco*RNAP holoenzyme synthesized a single 3-nt RNA (pppC[α-^32^P]UpG) starting 8 nt downstream of the 3’ end of the template DNA (designated as “+1”) (**Fig. 1B**). This start site assignment was validated by using the antibiotic rifampicin that inhibits the synthesis of the second phosphodiester bond. Indeed, rifampicin addition abolished the formation of 3-nt RNA and induced the accumulation of radiolabeled 2-nt RNA (pppC[α-^32^P]U) (**Fig. 1C**).

**Figure 1.**
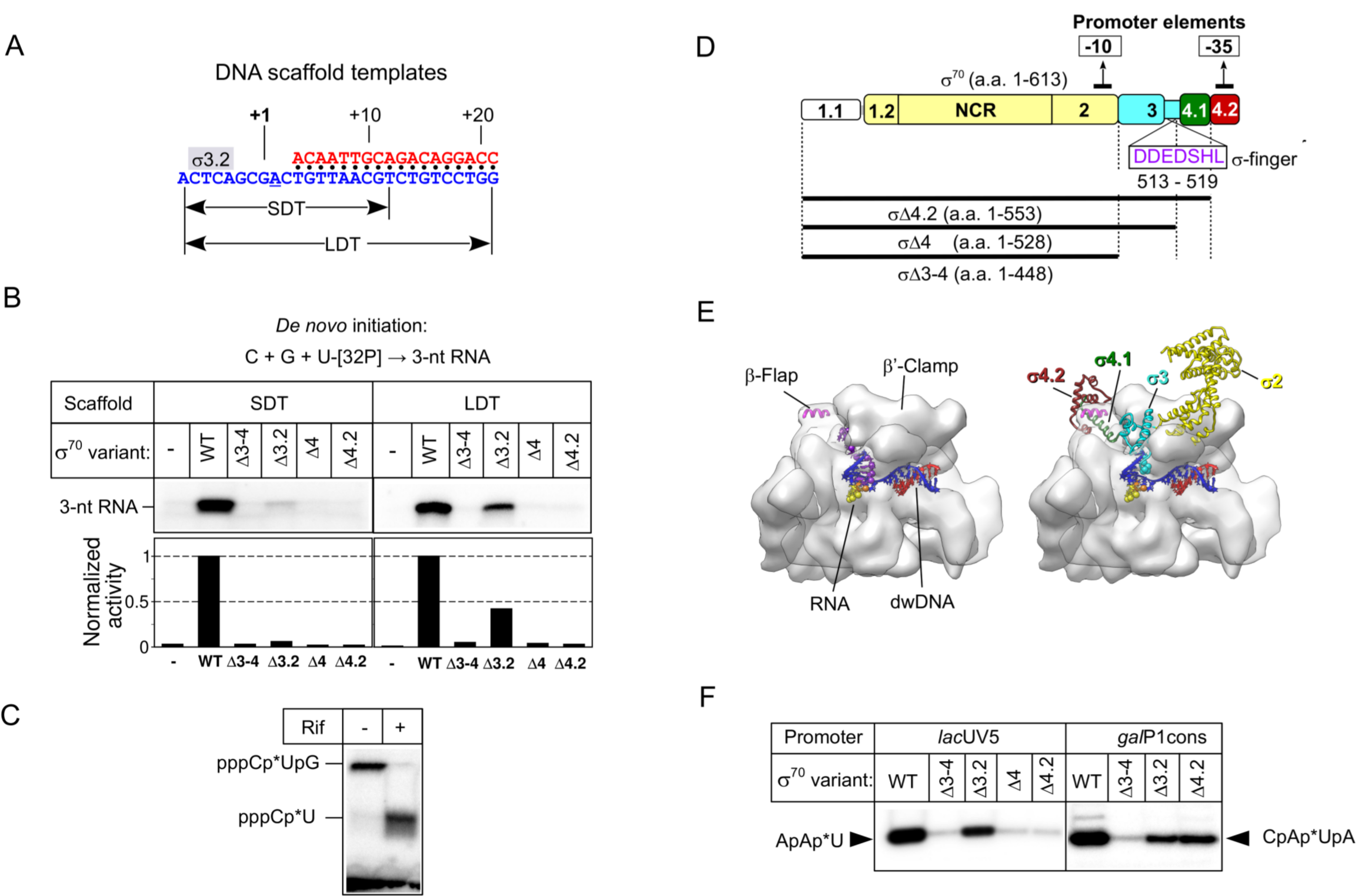
The σ subunit stimulates *de novo* transcription initiation on promoter-less DNA scaffolds. (A). Scheme of the synthetic DNA scaffolds: <u>S</u>hort <u>D</u>uplex <u>T</u>emplate (SDT) and <u>L</u>ong <u>D</u>uplex <u>T</u>emplate (LDT). Blue, template strand; red, non-template DNA strand. The position of the σ3.2-finger (σ3.2) is indicated by a gray rectangle. (B). Transcription initiation by wild type (WT) *Eco*RNAP, or harboring the indicated mutant σ^70^ variants, at the SDT and LDT scaffolds in the presence of CTP, GTP and [α-^32^P]-UTP. The bar graph shows the RNA product quantification. The RNA amount in each lane was normalized to the RNA synthesized in the presence of the full length σ^70^ subunit. (C). Inhibition of transcription on SDT by 10 μM rifampicin (Rif). (D). Scheme showing the σ^70^ subunit with domains 1 to 4. NCR, non-conserved region. The regions interacting with the promoter −10 and −35 consensus elements are indicated. The organization of the mutant σ^70^ variants is shown with black lines underneath the scheme. The σ3.2-finger sequence (amino acids 513-519) deleted in the σ_Δ3.2_ mutant is shown in purple letters. (E). Structural models of the *Eco*RNAP core (left) and holoenzyme (right) (PDB: 6C9Y) shown as semitransparent surfaces in complex with scaffold DNA (blue-red) (DNA and RNA from PDB: **2O5I**). The core is in complex with 14-nt RNA (yellow-orange-purple) and the holoenzyme is in complex with 4-nt RNA (yellow-orange). The RNA 5’-phosphates are shown as spheres. The 5’ end nucleotide is colored in orange. The σ^70^ subunit is shown as ribbons with the conserved regions colored as in **panel D**. The Cα atoms of amino acids 513-519 in region 3.2 are shown as spheres. (F). Transcriptional activity of the *Eco*RNAP holoenzyme harboring mutant σ^70^ variants. Transcription was initiated at the *lac*UV5 promoter by the ApA primer and [α-^32^P]-UTP and at the *gal*P1cons promoter by the CpA primer, [α-^32^P]-UTP and ATP. The RNA product sequences are indicated.

### The promoter-binding domain σ4 is essential for *de novo* initiation of RNA synthesis

To identify the σ^70^ regions that influence RNA synthesis initiation on minimal scaffolds, we generated a panel of σ^70^ mutants (**Fig. 1D, E**). The σ_Δ3-4_ fragment (residues 1-448) in which the regions 3 and 4 were deleted, is inactive in promoter-dependent transcription initiation (Zenkin *et al*, 2007). The σ_Δ4_ fragment (residues 1-529) lacked part of region 3.2 and the entire σ4 domain. In the σ_Δ4.2_ fragment (residues 1-553), only region 4.2 was deleted, but not the 4.1 α-helix, which binds inside the RNA exit channel (**Fig. 1E**). In agreement with previous studies (Kumar *et al*, 1993)(Kumar *et al*, 1994), the *Eco*RNAP holoenzymes harboring the σ_Δ4_ or σ_Δ4.2_ fragments were inactive in abortive transcription assays with the −10/−35 type *lac*UV5 promoter, and displayed reduced transcriptional activity with the “extended −10” type *gal*P1cons promoter (**Fig. 1F**). The σ_Δ3.2_ subunit, in which residues 513-519 in the σ3.2-finger were deleted, was active in transcription initiation with both promoters. The *Eco*RNAP holoenzymes assembled with the mutant σ^70^ subunits were inactive in *de novo* transcription initiation on the SDT scaffold (**Fig. 1B**). We detected no synthesis of dinucleotide RNA products by the *Eco*RNAP core and by the holoenzyme, differently from what reported for initial transcription on the M13 minus-strand origin (Zenkin & Severinov, 2004). This difference might be explained by the low NTP concentration (22 μM) used in our experiments. Conversely, on the LDT scaffold, the activity of σ_Δ3.2_ corresponded to 42% of the activity of full length σ^70^. Thus, strengthening the interaction between RNAP and the dwDNA duplex beyond position +10 can compensate for the lack of interaction between the σ3.2-finger and template-strand ssDNA. This result suggests that the σ3.2-finger/DNA interaction contributes to, but is not essential for initial transcription. The transcription defects caused by deletions in domain σ4, which does not interact with scaffold DNA, cannot be compensated by the dwDNA interactions, suggesting that σ4.2 is essential for initial transcription and exerts its activity through interaction with RNAP. Conversely, it has been suggested that σ4 is dispensable for initial transcription on the M13 phage minus-strand origin (Zenkin & Severinov, 2004). This discrepancy might be caused by differences in the DNA template architecture.

### dwDNA duplex and RNA primers suppress the translocation defect caused by deletions in the σ subunit

In promoter-dependent transcription initiation, short (≤ 3-nt) RNA primers (pRNAs) can rescue the defects linked to deletions in the σ3.2 and σ4 regions (Campbell *et al*, 2002)(Zenkin & Severinov, 2004) (Kulbachinskiy & Mustaev, 2006). To determine whether they have the same effect also when using minimal scaffold templates, we carried out transcription in the presence of a 2-nt pRNA (GpC, pRNA2) the 3’ end of which was complementary to the third position upstream of the DNA duplex (designated as position “+1”) (**Fig. 2A**). We assumed that the first catalytic step, addition of [α-^32^P]-UTP to pRNA2 (synthesis of GpC[α-^32^P]U), does not requires translocation of the RNAP active center because the 3’ end of pRNA2 binds to the “product-site” (i-site, facing the position +1 of the DNA template), thus leaving the “substrate site” (i+1, facing position +2 of the DNA template) available for incoming NTP (**Fig. 2B**). This hypothesis is supported by the structures of the RNAP initiation complexes with synthetic scaffolds observed in post-translocated states (Cheung *et al*, 2011)(Zhang *et al*, 2012). The next catalytic steps (synthesis of the GpC[α-^32^P]UpG and GpC[α-^32^P]UpGpA products) require the translocation of the RNA 3’ end from the i+1 site to the i-site (**Fig. 2A, B**). As only the first incorporated NTP (U) was labeled, the fraction of the longest reaction product (RNA[N+2]) reflected the overall “efficiency of RNAP translocation” from register +2 to + 4.

**Figure 2.**
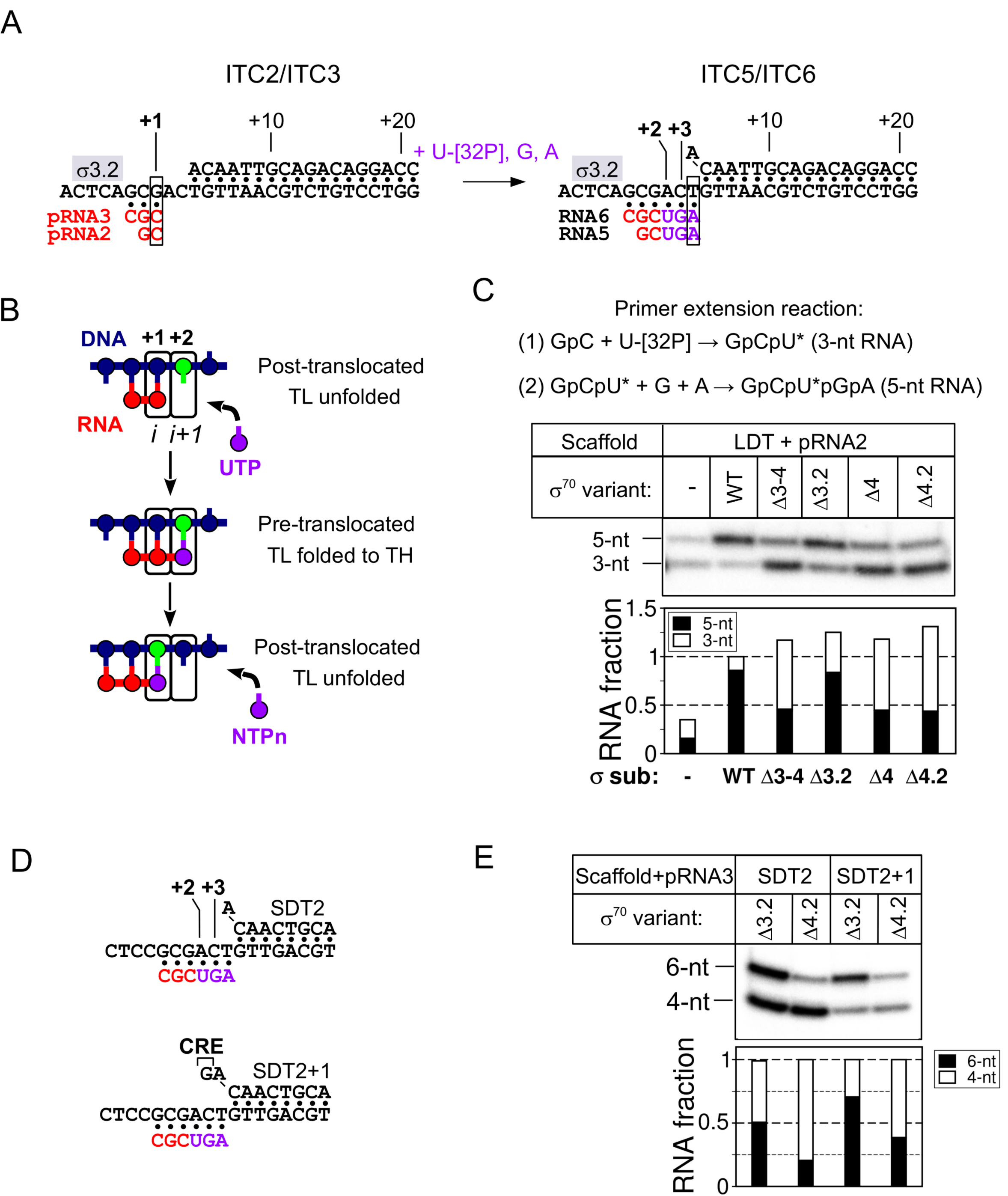
The σ subunit and DNA template architecture modulate forward translocation. (A). Scheme of the primer extension reaction on the LDT scaffold. The RNA primers (pRNA) used to assemble Initial <u>T</u>ranscribing <u>C</u>omplexes (ITCs) are shown in red. Nucleotides added during initial transcription are in purple. The position of the σ3.2-finger is indicated by a gray rectangle. (B). Simplified scheme of the nucleotide addition cycle (NAC) on scaffold DNA template (blue) with a 2-nt RNA primer (red) and initiating 3’ UTP (purple). The active site registers are designated as i and i+1. TL, trigger loop; TH, trigger helix. (C). Extension of 2-nt pRNA (pRNA2) by *Eco*RNAP on the LDT scaffold in the presence of wild type (WT) or mutant σ^70^ variants. The stacked bar graph shows the RNA product quantification. The RNA amount in each lane was normalized to the total RNA synthesized in the presence of the full length σ^70^ subunit. (D). Scheme of the SDT2 and SDT2+1 scaffolds with 6-nt nascent RNA. Red, sequence of the 3-nt pRNA; purple, nucleotides added during primer extension. (E). Transcription initiation by *Eco*RNAP at the SDT2 and SDT2+1 scaffolds with 3-nt pRNA (pRNA3) in the presence of mutant σ^70^ variants. The stacked bar graph shows the RNA product quantification. The amounts of 4-nt and 6-nt RNA in each lane were normalized to the total RNA (4-nt + 6-nt) amount in each lane.

In the presence of GpC and three nucleotides ([α-^32^P]-UTP, GTP, and ATP), the *Eco*RNAP core synthesized two [^32^P]-labeled RNA products (3-nt and 5-nt RNAs), with a bias toward the shorter one (overall translocation efficiency: ∼ 40%, **Fig. 2C**). In the same conditions, σ^70^ strongly stimulated [α-^32^P]-UTP incorporation and translocation. Consequently, the 5-nt RNA was the major reaction product synthesized by the *Eco*RNAP holoenzyme (95% translocation efficiency). The observed low efficiency of the [α-^32^P]-UTP-addition reaction by the *Eco*RNAP core might reflect its low affinity for pRNA2. Indeed, increasing pRNA2 concentration increased [α-^32^P]-UTP incorporation, but did not stimulate translocation (**Fig. S1**). Even the “minimal” σ_Δ3-4_ fragment strongly stimulated [α-^32^P]-UTP incorporation, compared with the *Eco*RNAP core (**Fig. 2D, Fig. S2**). Therefore, binding of domain σ2 to the *Eco*RNAP core may strengthen its interaction with the scaffold DNA/RNA and/or directs it to the active site cleft.

Like with promoter DNA templates, the initiation defect on DNA scaffolds upon σ3.2-finger deletion was rescued by pRNA2 (∼80% translocation efficiency, **Fig. 2C**). The C-terminal deletions in the σ^70^ subunit (σ_Δ3-4_, σ_Δ4_ and σ_Δ4.2_) hindered the synthesis of 5-nt RNA (<50% translocation efficiency) leading to RNA patterns similar to those of the *Eco*RNAP core. As deletion of the region 4.2 and deletion of the entire σ3 and σ4 domains led to the same effect, we conclude that σ4.2 is a principal determinant for the efficient synthesis of the 5-nt RNA product. It is unlikely that the translocation defect conferred by deletion of the region σ4.2 is due to a decreased affinity of the 3-nt RNA product (GpC[α-^32^P]U) for the RNAP active site because the overall [α-^32^P]-UTP incorporation was similar with full length σ^70^ and σ_Δ4.2_ (**Fig. 2C**). Our results suggest that RNAP translocation from register +2 to register +3 is a rate-limiting step for 5-nt RNA synthesis, and is slower than [α-^32^P]-UTP addition and the 3-nt RNA product dissociation. The σ4 deletions had similar effects on translocation also when using the SDT scaffold and the 3-nt pRNA (CpGpC, pRNA3) (**Fig. S2**). However, the 3-nt pRNA suppressed the transcription defects caused by the σ4 deletion on the LDT but not the SDT scaffold, indicating the RNAP interaction with dwDNA stimulates translocation. To explore the effect of the DNA duplex on RNAP translocation we used modified versions of the SDT scaffold (SDT2 and SDT2+1), with more stable dwDNA duplexes (**Fig. 2D**). Moreover, in the SDT2+1 scaffold, the 5’ end upstream edge of the DNA duplex was extended by one base pair (G:C). Thus, translocation from register +2 to register +3 on SDT2+1 requires the unpairing of 1 bp of dwDNA followed by the formation of the contact between G at position +3 of the non-template DNA and the CRE pocket of the RNAP core that is known to counteract transcriptional pausing (Vvedenskaya *et al*, 2014). Translocation was more efficient on the SDT2+1 scaffold compared with the SDT1 scaffold (**Fig. 2E**), suggesting that dwDNA duplex melting is not a barrier for translocation and that the interaction with CRE stimulates translocation. We conclude that *Eco*RNAP pauses after the addition of the first NTP to the RNA primer and that the interaction with the downstream DNA and RNA promotes forward translocation. The region σ4.2 may act on translocation by strengthening this interaction.

### The σ subunit regions 3.2 and 4.2 stabilize ≤ 4-nt RNAs in the RNAP active site

To determine whether the σ subunit can stabilize short RNAs in the RNAP active site, we immobilized ITCs on Ni^2+^-agarose beads and tested their ability to retain RNAs by washing the complexes with transcription buffer. We used 2 to 6 nt-long pRNAs the 3’ end of which was aligned to the same position of the template, designated as position “+1” (**Fig. 3A**) Control experiments in which complexes were formed by the core *Eco*RNAP on SDT and LDT scaffolds showed that after washing with the “high salt” buffer containing 1M NaCl (**Fig. S3**), 5-, 6-, 7-, 13-nt pRNAs were stably bound in ITCs. However, reduced retention of 5- and 6-nt pRNAs, was observed with the SDT scaffold. This indicates that the dwDNA duplex contributes to the overall stabilization of the complex. Consequently, the LDT scaffold was used in the next experiments. To measure the retention efficiency, ITCs containing 2-to 6-nt RNAs were either washed with the transcription buffer containing 250 mM NaCl and then labeled with [α-^32^P]-UTP, or directly labeled without washing step (**Fig. 3B**). This experiment demonstrated that 2-to 4-nt pRNAs were weakly bound to ITCs compared with 5-to 6-nt pRNAs (**Fig. 3B-D**). Therefore, the 4-bp RNA:DNA hybrid is a conversion point between stable and unstable ITC states. Moreover, we observed a clear difference in the capacity to hold 4-nt pRNA by ITCs containing the full length σ subunit (75% retention efficiency) and σ mutants (σ_Δ3.2_ and σ_Δ4_; <50% retention efficiency) (**Fig. 4E)**. The defect was stronger for σ_Δ4_-*Eco*RNAP than σ_Δ3.2_-*Eco*RNAP. We obtained similar result with the SDT scaffold (**Fig. S4**). To determine whether the slight difference in 4-nt pRNA retention observed between σ_Δ3.2_ and σ_Δ4_ (**Fig. 4E)** was significant, we performed several washing steps on ITCs formed by σ_Δ3.2_-*Eco*RNAP and σ_Δ4.2_-*Eco*RNAP (**Fig. 4F**,**G**). After the third washing step, almost no bound RNA was left in σ_Δ4.2_-ITC (∼10% retention relative to full length σ^70^), while RNA retention was higher for σ_Δ3.2_-ITC (∼40% retention relative to full length σ^70^). RNA binding remained stable with wild type σ^70^-ITC (60% retention relative to the ‘no washing’ condition). We conclude that the σ^70^ subunit stabilizes 4-5-nt-long RNAs in the RNAP active site and that the region σ4.2 is a major determinant of this activity.

**Figure 3.**
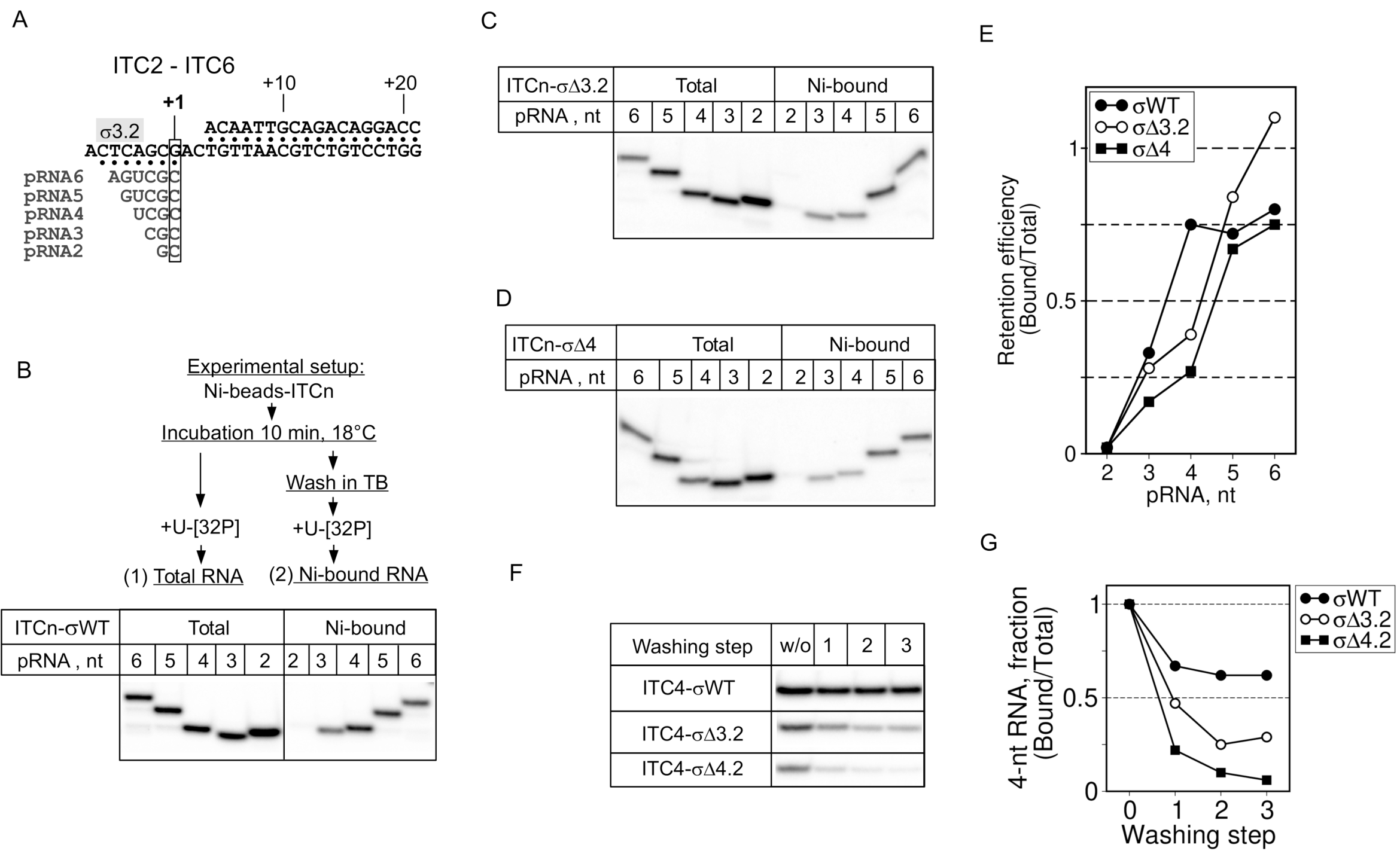
The σ subunit regions 3.2 and 4 stabilize short RNAs in ITCs. (A). Scheme of the LDT scaffold with the RNA primers (pRNA) used to assemble ITC2 - ITC6. (B, C, D). [α-^32^P]-UTP-labeled RNAs produced in primer extension reactions in the presence of σ^70^, σΔ3.2, and σΔ4, respectively. ITCs were immobilized on Ni^2+^-agarose beads and washed (Ni-bound) or not (Total) with buffer (TB), as indicated, before labeling. The experimental setup is shown schematically on top of panel B. (E). Quantification of the results shown in panels B,C,D. The fraction of labeled RNA retained in ITCn was calculated as the ratio between the RNA amount in the washed ITCn (Ni-bound) to the RNA amount in the unwashed ITCn (Total). (F). Comparison of the retention efficiency for 4-nt RNA in the presence of σ^70^ (WT), σΔ3.2 and σΔ4.2. The number of washing steps is indicated; w/o, without washes. (G). Quantification of the results shown in panel F. The fraction of labeled RNA retained in ITC4 is plotted as a function of the washing step number. The RNA amount in each lane was normalized to the RNA in unwashed ITC4.

**Figure 4.**
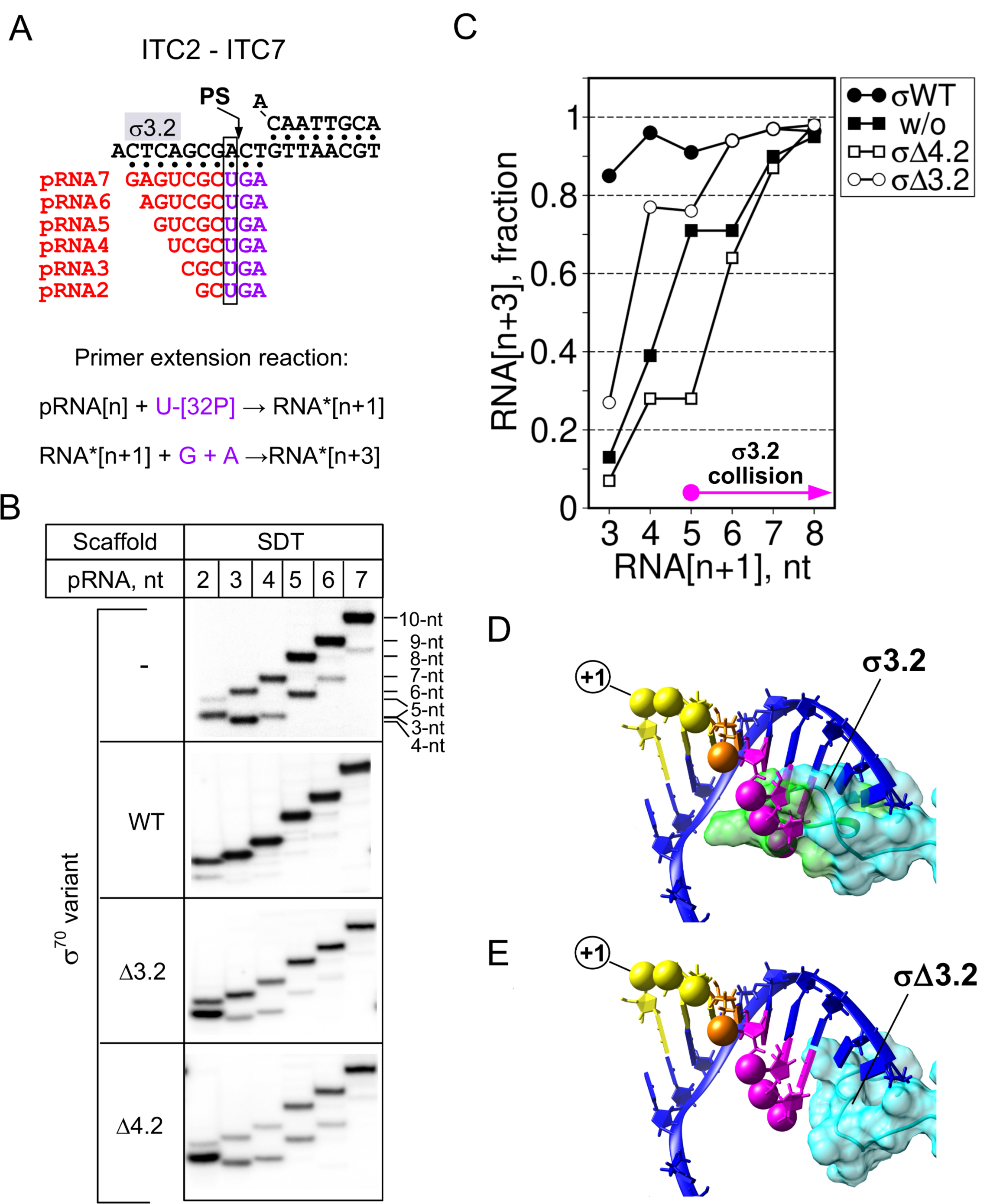
RNA:DNA hybrid and σ modulate RNAP translocation efficiency. (A). Scheme of the SDT scaffold with primer RNAs (pRNA) of various lengths (red) used to assemble ITC2 - ITC7. Nucleotides added during initial transcription are in purple. (B). Primer extension reactions, shown schematically at the top, were performed with the pRNAs shown in panel A in the presence or absence of the indicated σ^70^ variants. (C). Quantification of the experiments shown in panel B. The values of RNA[n+3] for each lane were normalized to the total RNA (RNA[n+1] +RNA[n+3]) in each lane and were plotted as a function of the RNA length. (D, E). Structural models of the σ^70^ subunit-DNA-RNA interactions in the RNAP main channel. The σ3.2 finger (PDB: **4YG2**) is shown as a molecular surface. The coordinates of the DNA template strand (in blue) and RNA are from PDB: **2O5I**. RNA 5’-phosphates are shown as spheres colored in function of the RNA length. The RNA 3’ end is in yellow, the RNA 5’ end nucleotides that clash with σ3.2-finger are in magenta. The nucleotide in the fourth position, marking the transition from unstable to stable ITC, is in orange. (E). The same as in panel D, with the σ3.2-finger residues (aa 513-519) deleted. Models were built with COOT (Emsley & Cowtan, 2004) and UCSF Chimera (Pettersen *et al*, 2004).

### The σ region 4.2 promotes extension of ≤ 7 nt-long RNAs

To explore the relationship between translocation efficiency and RNA:DNA hybrid stability, we performed 2-min primer extension reactions with pRNAs of various lengths in the presence of [α-^32^P]-UTP, GTP and ATP (**Fig. 4A**). In these experiments, to facilitate the detection of the initial pause, we used the SDT scaffold that displayed stronger σ-dependence in translocation. The *Eco*RNAP core efficiently extended ≥ 8-nt pRNAs (>90% efficiency) (**Fig. 4B**,**C**), and its translocation efficiency decreased gradually with the RNA length shortening. This dependence on RNA length might be explained by the intrinsic instability of RNA:DNA hybrids and/or by the disengagement of the RNA 3’ end from the active site. The translocation efficiency was independent from the RNA length when the wild type *Eco*RNAP holoenzyme was used (**Fig. 4B,C**). The σ_Δ3.2_-*Eco*RNAP holoenzyme displayed strong translocation defects with 3-nt pRNA (∼25% efficiency), moderate defects with 4-5-nt pRNAs (∼80% efficiency), and no defect with 6-nt pRNA (>90% efficiency). On the basis of the RNAP-promoter complex structure data, the 5’ end of ≥ 5-nt-long RNAs clashes with the σ3.2-finger (**Fig. 4D**). Thus, the σ3.2-finger should hinder the extension of RNAs longer than 5 nt and favor abortive initiation (Murakami *et al*, 2002)(Li *et al*, 2020). A model of the σ3.2-finger deletion on the *Eco*RNAP structure (**Fig. 4E**) showed that 5-to 7-nt RNAs can be accommodated in the active site cleft. Yet, the remaining segment of the region σ3.2 can still contact the template DNA strand at positions −6/-7. Strikingly, in our experiments, ‘abortive’ RNAs accumulated when using the *Eco*RNAP core and the σ_Δ3.2_-*Eco*RNAP holoenzyme, but not with the wild type *Eco*RNAP holoenzyme. This suggests that the σ3.2-finger is not the primary cause of abortive RNA formation and that abortive RNA synthesis is not an obligatory event in initiation (Duchi et al. 2016).

Unlike σ_Δ3.2_-*Eco*RNAP, translocation stimulation was defective with the σ_Δ4.2_-*Eco*RNAP holoenzyme and pRNAs shorter than 8 nt. The properties of the σ_Δ4.2_-*Eco*RNAP holoenzyme were identical to those of the *Eco*RNAP core except with 5-nt-long RNA that displayed increased translocation efficiency with the *Eco*RNAP core. We did not investigate the reason of this unusual behavior. The results of the “RNA-retention” experiments in combination with the “translocation-dependence” experiments demonstrated that there is no correlation between RNA:DNA hybrid stability and translocation efficiency. Indeed, 5- and 6-nt RNAs were stably bound to ITC, but still displayed σ-dependence for translocation. Thus, we conclude that the low efficiency in nascent RNA extension (any lengths) by the σ_Δ4.2_-*Eco*RNAP holoenzyme is due to RNAP pausing after the first NTP addition, and that the stimulation of RNAP translocation by σ is unlikely to occur through RNA:DNA hybrid stabilization.

### The RNA 3’ end nucleotide identity determines the initial-transcription pause duration

To explore the impact of σ4.2 on pausing, we studied the kinetics of 5-nt and 7-nt RNA synthesis initiated with pRNA3 (ITC3, unstable RNA:DNA hybrids) and pRNA5 (ITC5, stable RNA:DNA hybrids), respectively **(Fig. 5A)**. The overall [α-^32^P]-UTP addition rate was similar for ITC3 and ITC5 in the presence of the σ_Δ4.2_, and was highest in the presence of the full length σ^70^ (**Fig. 5B**,**C**). Translocation from register +4 to +5 was at least 100-fold faster (t_1/2_ ∼ 1.7 s) in the presence of full length σ^70^ than of the σ_Δ4.2_ mutant (t_1/2_ ∼ 200 s) (**Fig. 5B,D**). Reactions were completed in 120 s, without any further incorporation of [α-^32^P]-UTP. Therefore, the labeled 5-nt RNA product remained bound to RNAP. Extension of the pRNA 5’ end by 2 nucleotides (**pRNA5)** accelerated the forward translocation only by 2-fold. Therefore, in agreement with the conclusion drawn in the previous section, the overall RNA:DNA hybrid stabilization has little effect on pausing.

**Figure 5.**
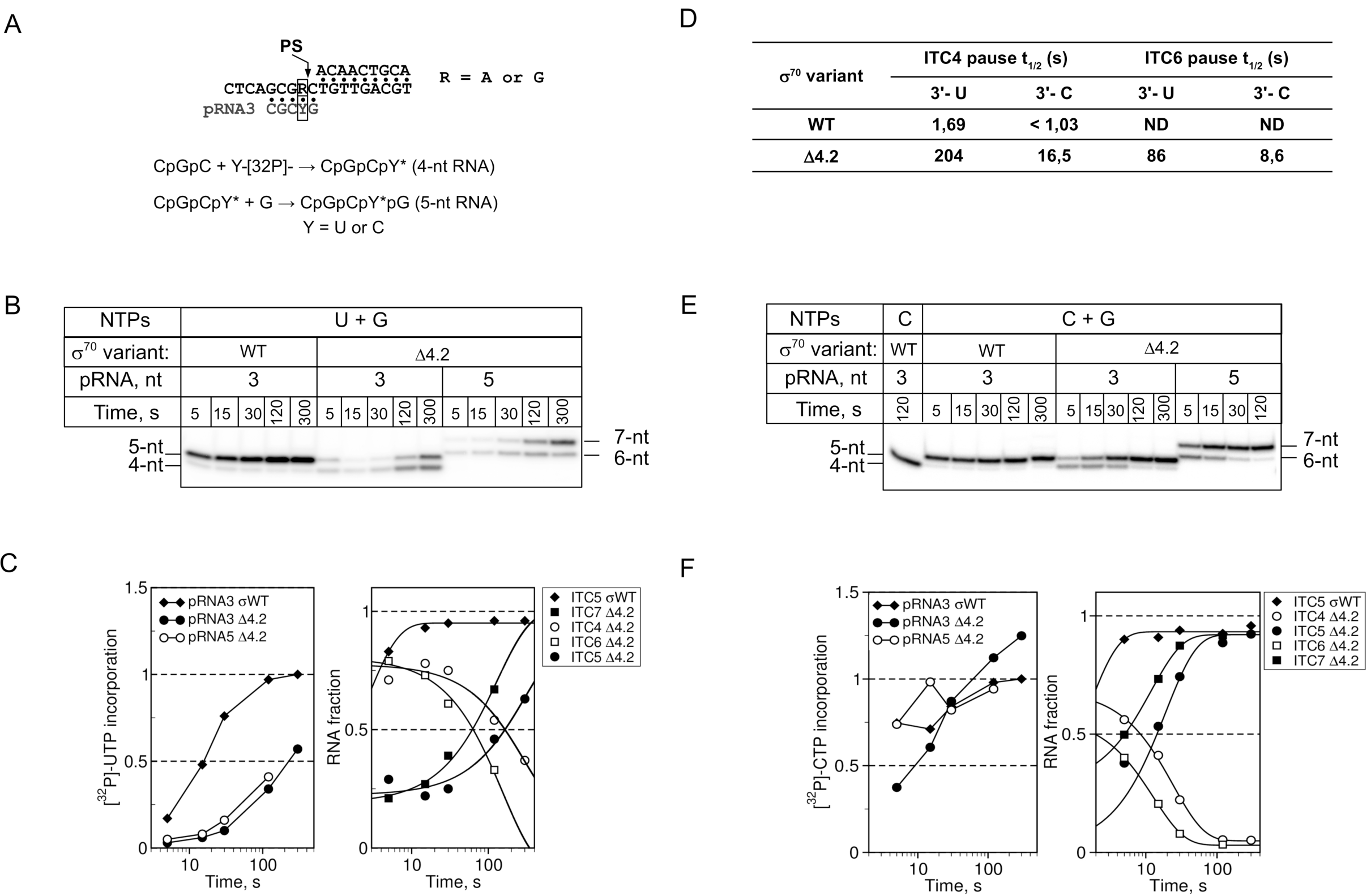
The RNA 3’ end nucleotide modulates the initial pause duration. (A). Scheme of SDT2 and SDT-G scaffolds with 3-nt primer RNA (pRNA3). The primer extension reactions were performed with [α-^32^P]-UTP on the SDT scaffold and with [α-^32^P]-CTP on the SDT-G scaffold. (B). Kinetics of primer extension at the SDT scaffold. **(C)**. Quantification of the experiment shown in panel B. The left graph shows the synthesis of total RNA over time. The right graph shows the RNA product fractions, representing the different ITCs, over time. For each time point, values were normalized to the total RNA synthesized at that time point. **(D)**. Pause half-life times (t_1/2_) calculated from the experiments shown in panels B and E. **(E)**. Kinetics of primer extension on the SDT-G scaffold. **(F)**. Quantification of the experiment shown in panel E.

In our assay, the RNA chain elongation starts with addition of U that forms an unstable U:dA pair with the DNA template (Huang *et al*, 2009). Therefore, the 3’ end nucleotide might disengage from the active site, and induce pausing. If this hypothesis is correct, the substitution of the U:dA pair by the more stable C:dG pair should suppress pausing, and favor forward translocation. To test this assumption, we used a scaffold (SDT-G) harboring G instead of A at position +2 (**Fig. 5A**), and initiated the primer extension with [α-^32^P]-CTP. Unlike UTP, the CTP addition rate with the σ_Δ4.2_**-***Eco*RNAP holoenzyme was close to that observed with the wild type *Eco*RNAP holoenzyme (compare **Fig. 5C and 5F**). Thus, without σ4, *Eco*RNAP senses the difference between UTP and CTP, while CTP suppresses the effect of the σ4 deletion. Irrespectively of the RNA length (4-nt or 6-nt), translocation of the mutant σ_Δ4.2_**-***Eco*RNAP from the register +2 to the register +3 was accelerated by ∼10-fold on SDT-G DNA compared with SDT-A DNA. This suggests that the stability of base pairing at the 3’ end, but not the RNA:DNA hybrid length is crucial for forward translocation. As σ_Δ4.2_**-***Eco*RNAP translocation rate was significantly reduced even when the RNA 3’ end was stabilized, compared with wild type *Eco*RNAP, we conclude that the region σ4.2 may affect the active site cycling or the clamp opening-closing dynamics that control RNAP translocation.

### The σ4 remodeling co-activator AsiA stimulates pausing

Region σ4.2 binds to the flap-tip-helix (FTH) of the RNAP β subunit (Kuznedelov *et al*, 2002) (Geszvain *et al*, 2004) that is implicated in the regulation of pausing (Kang *et al*, 2018). To test whether σ4.2 exerts its effect on RNA synthesis through interaction with β-FTH, we used the T4 phage co-activator protein AsiA. AsiA remodels exactly the same region in the σ^70^ subunit (residues 528 - 613) that was deleted in the σ_Δ4_ mutant, and disrupts the interaction between σ4.2 and β-FTH (Hinton & Vuthoori, 2000) (Shi *et al*, 2019) (model in **Fig. 6A**). If the σ4.2-β-FTH contact were essential for RNA synthesis, AsiA should fully inhibit initial transcription. To test this hypothesis, we performed *de novo* and primed transcription by the σ^70^-*Eco*RNAP and σΔ4.2-*Eco*RNAP holoenzymes, with and without AsiA, on the SDT2 template (**Fig. 6B**). As control, we used an abortive transcription assay on the *lac*UV5 promoter. AsiA inhibited *lac*UV5-dependent transcription initiation by 85% (**Fig. 6B, lanes 1**,**2 and Fig. 6C**). Conversely, *de novo* initiation from the scaffold was much less sensitive to AsiA. Indeed, 3-nt RNA synthesis was inhibited only by 50%, which coincided with the accumulation of the short 2-nt RNA product (**Fig. 6B, lanes 7**,**8 and Fig. 6C**). AsiA also influenced transcription initiated with the GpC primer (**Fig. 6B, lanes 3**,**4 and Fig. 6C**). The amount of 3-nt RNA increased simultaneously with the increase in total [α-^32^P]-UTP incorporation. Such effect was consistent with the AsiA-induced destabilization of short RNAs, leading to accumulation of “abortive” transcripts. The finding that AsiA did not affect [α-^32^P]-UTP incorporation with the σΔ4-*Eco*RNAP holoenzyme (**Fig. 6B, lanes 5**,**6 and Fig. 6C**) indicates that AsiA modulates RNA synthesis through σ4. However, the weak impact of AsiA on initial transcription was in striking contrast with the strong inhibitory effect of the σ4 deletion. The only possible explanation for this discrepancy can be that the σ4 physical presence in the RNA exit channel is essential for initial transcription. In the presence of AsiA, the σ4 domain remains bound inside the RNA exit channel (Shi *et al*, 2019), and therefore AsiA exerts only a weak effect on scaffold-dependent transcription. We conclude that the interaction of σ4.2 with β-FTH modulates the catalytic site activity, but is not essential for initial transcription.

**Figure 6.**
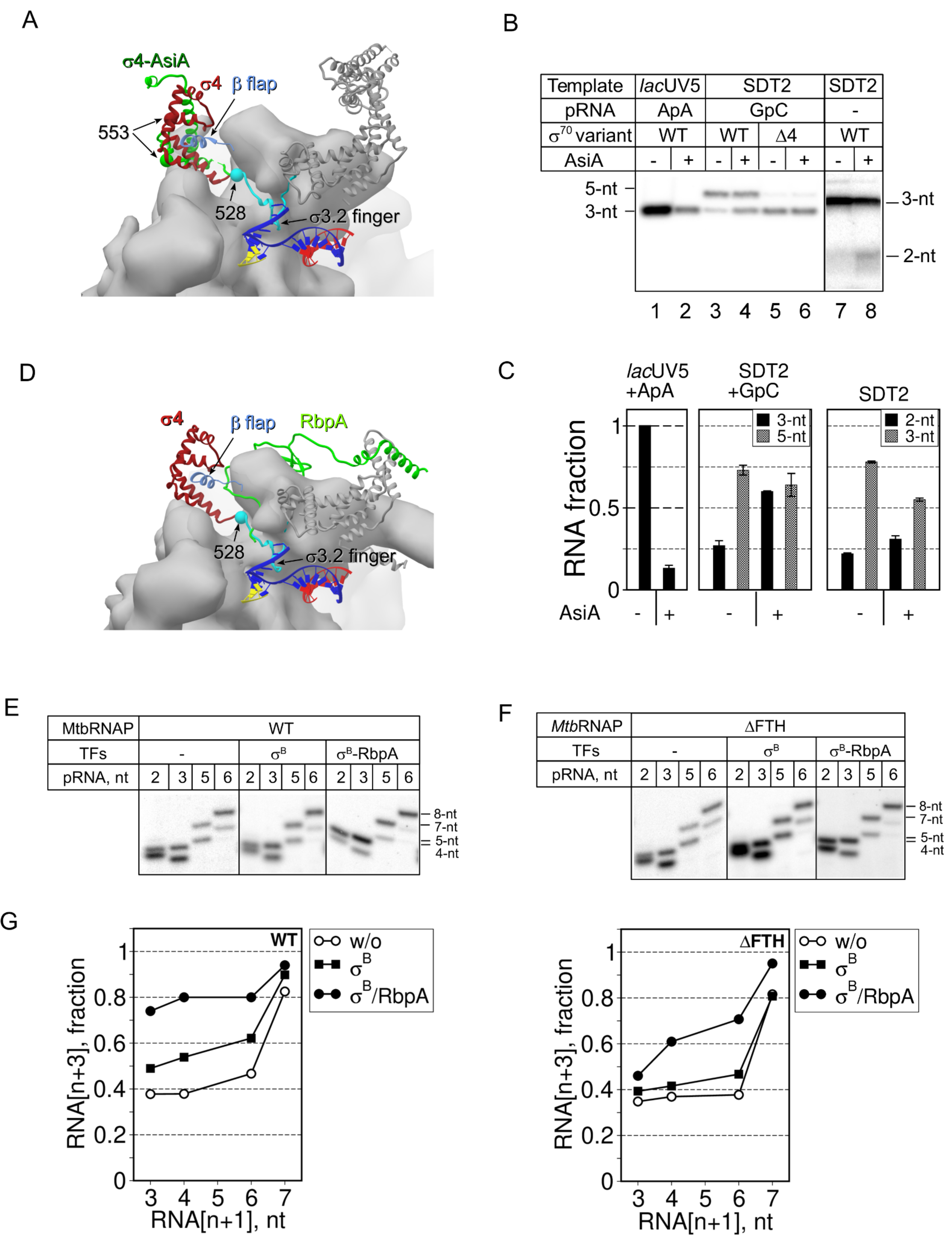
The σ-remodeling factors AsiA and RbpA modulate initial transcription. (**A**). Structural model of the σ^70^-*Eco*RNAP holoenzyme (PDB: **4YG2**) in complex with the SDT scaffold DNA and 3-nt RNA (yellow). Core RNA is shown as molecular surfaces. The β subunit is shown as a transparent surface. The σ subunit is shown as a ribbon in which the σ4 domain is in dark red, the σ region 3.2 is in cyan, and the other regions are in gray. The β-Flap (G891-G907) is shown as a ribbon (cornflower blue). The remodeled conformation of the σ4 domain (σ4-AsiA) in the *Eco*RNAP-AsiA-MotA complex (PDB: **6K4Y**) is shown in light green. The Cα atoms of the σ^70^ residues 528 and 553, shown as spheres, mark the borders of the σΔ4 and σΔ4.2 deletions, respectively. (**B**). Transcription initiation at the *lac*UV5 promoter and SDT2 scaffold in the presence or not of AsiA. Transcription was initiated by ApA and [α-^32^P]-UTP (lanes 1,2), by the GpC primer, GTP and [α-^32^P]-UTP (lanes 3-6), or by CTP, GTP and [α-^32^P]-UTP (lanes 7,8). (**C**). Quantification of the experiments shown in panel B. For each experimental condition (-/+ AsiA), the amount of each RNA product was normalized to the total RNA synthesized without AsiA. Averages and standard errors of two independent experiments are shown. (**D**) Structural model of σ^A^-*Mtb*RNAP (PDB: **6EDT**) in complex with RbpA (green ribbon) and the SDT scaffold DNA. The nomenclature and color code are the same as in panel A. (**E**) Primer extension reactions performed by the *Mtb*RNAP core and σ^B^-*Mtb*RNAP holoenzyme on the SDT2 scaffold with and without RbpA. Transcription was initiated by GTP and [α-^32^P]-UTP. (**F**) Primer extension reactions performed by the *Mtb*RNAP^ΔFTH^ core and σ^B^-*Mtb*RNAP^ΔFTH^ holoenzyme on the SDT2 scaffold with and without RbpA. Transcription was initiated by GTP and [α-^32^P]-UTP. (**G**). Quantification of the results shown in panels E and F, performed as in Figure 4C.

### RbpA from *M. tuberculosis* stimulates translocation through the σ4.2/β-FTH interaction

To determine whether the σ subunit anti-pausing activity can be observed with RNAP from other bacteria, we studied initial transcription by *Mtb*RNAP. We used the *M. tuberculosis* σ^B^ subunit that requires the activator protein RbpA to stabilize its active conformation in the *Mtb*RNAP holoenzyme (Vishwakarma *et al*, 2018). As RbpA N-terminus binds within the RNA-exit channel, it could modulate σ4.2 anti-pausing activity (model in **Fig. 6D**). First, we compared the kinetics of 5-nt pRNA extension by *Mtb*RNAP and *Eco*RNAP on SDT2 DNA in the presence of [α-^32^P]-UTP and GTP (**Figure S5**). As observed with *Eco*RNAP, *Mtb*RNAP paused at the register +6 during initiation from pRNA5, and its translocation was stimulated by the σ^B^-RbpA complex. To better understand the role of σ4.2-β-FTH interaction in initial pausing, we constructed a *Mtb*RNAP mutant in which the β subunit residues 811-825 were deleted (*Mtb*RNAP^ΔFTH^), and then assessed how the translocation activity of the mutant and wild type enzymes were influenced by the pRNA length (**Fig. 6E**,**F**). ITCs were assembled with 2, 3, 5 and 6-nt pRNAs (ITC2 to 6) in the presence of σ^B^ and RbpA, or without transcription factors, and supplemented with [α-^32^P]-UTP and GTP. As observed with *E*coRNAP, *Mtb*RNAP translocation efficiency increased gradually with the RNA length, and reached 80% with the 6-nt pRNA. Like for σ^70^, the σ^B^-RbpA complex stimulated the forward translocation with short pRNAs (3-6-nt in length). However, the σ^B^ subunit alone did not stimulate translocation, in agreement with fact that its conformation in the *Mtb*RNAP holoenzyme differs from that of σ^70^ in the *Eco*RNAP holoenzyme. The deletion of β-FTH abolished σ^B^ anti-pausing activity with ITC2 and ITC3 (2- and 3-nt pRNAs, respectively). Furthermore, the RNA amount produced by unstable ITC2/ ITC3 formed by the *Mtb*RNAP^ΔFTH^ mutant increased by ∼4-fold, compared with the amount produced by the stable ITC6. (**Fig. S6**). This “abortive-like” behavior was observed only in the presence of the σ^B^ subunit, and might be caused by a clash between the inappropriately positioned region 3.2 and RNA. Addition of RbpA only partially restored σ^B^ capacity to stimulate *Mtb*RNAP^ΔFTH^ translocation (**Fig. 6F and Fig. S5**). In agreement with the conclusion drawn from the experiments with AsiA, this result suggests that σ4.2 interaction with β-FTH regulates, but is not essential for the anti-pausing activity of σ^B^. The β-FTH deletion should dramatically destabilize σ4 positioning/interaction within RNA channel and consequently enhance pausing and abortive transcription. RbpA compensates for the lack of β-FTH probably by facilitating σ4 positioning within the RNA-exit channel.

## DISCUSSION

Our study demonstrates that RNAPs from different bacterial species, are inefficient in initial transcription and prone to pause upon extension of short RNAs (3 to 7-nt in length). The σ subunit region 4.2, which was implicated only in promoter binding, counteracts the initial transcription pausing and thus plays an essential role in organizing of the RNAP active center for efficient initiation of *de novo* RNA synthesis. The σ4.2 region displays two distinct activities: RNA:DNA hybrid-stabilizing activity, and translocation-stimulating activity. Modulation of these activities by the σ-remodeling factors may be a general mechanism to tune initial transcription.

### Initial transcription pausing on the pathway to abortive transcription

Our results suggest that at each nucleotide addition step, ITCs harboring 3-8 nt RNAs can enter into a paused state in which the RNA 3’ end is disengaged from the active site. The paused ITC (PITC) bifurcates in two pathways: abortive pathway in which nascent RNA dissociates from RNAP, and productive pathway in which nascent RNA remains bound to RNAP and slowly translocates to the next register (**Figure 7)**). PITC conversion to productive ITC is accelerated by (1) strengthening the dwDNA/RNAP interactions, (2) strengthening the RNA:DNA hybrid /RNAP interactions, and (3) stabilizing base pairing at the RNA 3’ end. These observations can be explained by a simple model in which the lack of the stable 9-bp RNA:DNA hybrid impedes the concerted translocation of RNA and DNA. During NAC, the RNAP pincers formed by the β’ clamp and β lobe should transiently adopt an open or partially open conformation (Vassylyev *et al*, 2007b)(Weixlbaumer *et al*, 2013). We speculate that this opening may weaken the RNAP interaction to hold together template DNA and RNA. Due to the thermal motions and the altered, misaligned structure of short RNA:DNA hybrids (Liu *et al*, 2011) (Cheung et al, 2011), the RNA 3’ end may be displaced from the active site. In addition, the 3’-U forms unstable base pairs (U:dA) that in the absence of σ4, may favor the formation of the frayed state, leading to backtracking and pausing (Artsimovitch & Landick, 2000) (Toulokhonov *et al*, 2007a). Consequently, the PITC remains blocked in one of the inactive states (half-translocated, hyper-translocated, or backtracked) that slowly isomerize to an active post-translocated state. The 3’-C that forms more stable (C:dG) base pairs remains in the active site, thus promoting forward translocation. In support to this model, the PITC half-life time was more strongly biased by the RNA 3’ end nucleotide identity than by the RNA:DNA hybrid length. Indeed, the RNA 3’ end nucleotide also modulated the translocation bias in stable elongation complexes with 8-9-bp RNA:DNA hybrids (Hein et. al, 2011). The σ subunit may restrain RNA and DNA motions by strengthening the RNAP core interactions with RNA and DNA, thus allowing the correct alignment of the template to the active site and promoting translocation. The σ-mediated stabilization of short RNAs in the active site and stimulation of the forward translocation should drastically reduce the probability of abortive transcription and shift the equilibrium toward promoter escape. As the half-life time of the initial pause strongly depends on the RNA 3’ end nucleotide identity, it should be a promoter-specific event determined by the initial transcribing DNA sequence.

**Figure 7.**
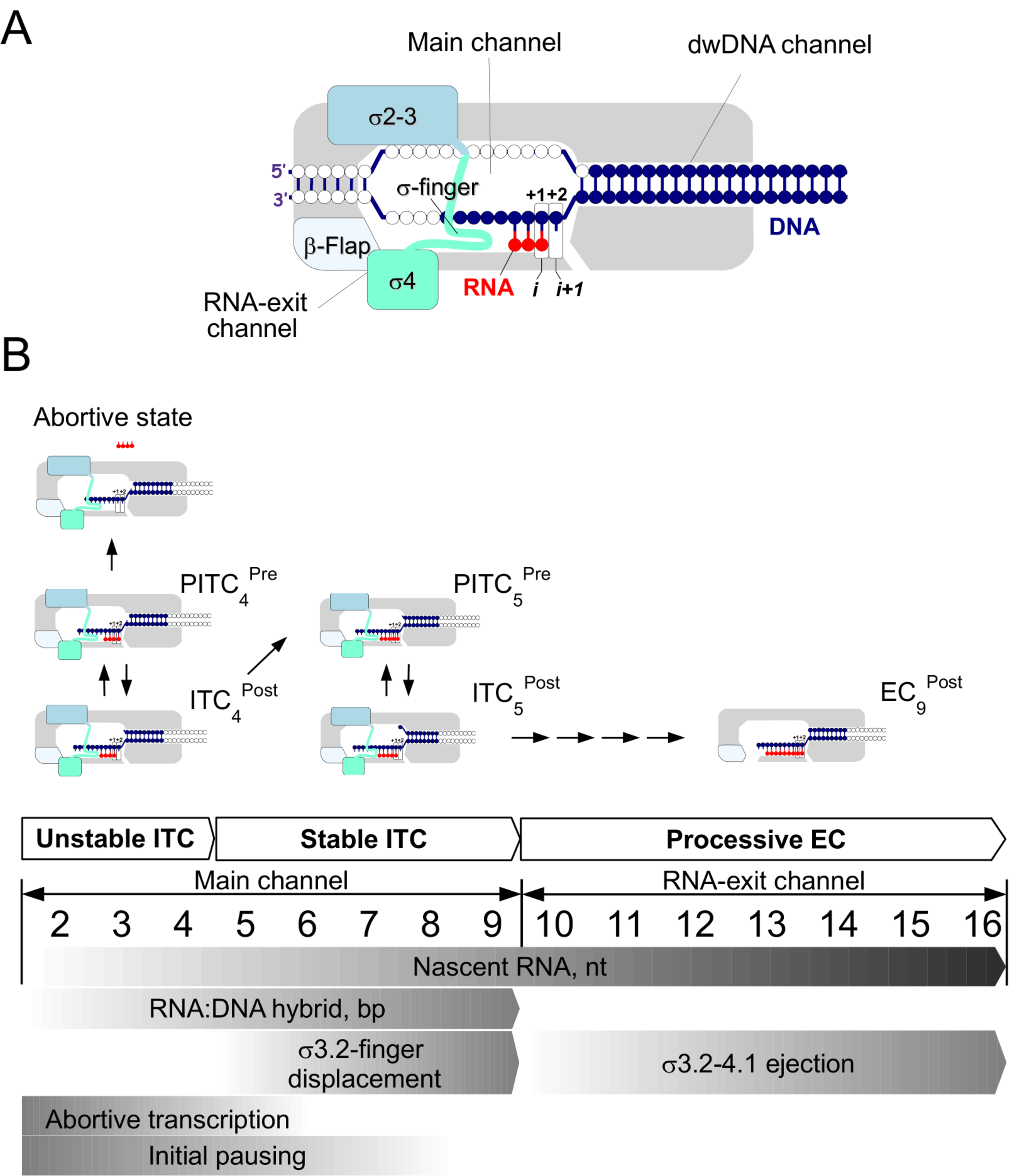
Model of events during initial transcription. (A). Schematic representation of RNAP in complex with DNA template. Bases corresponding to the scaffold DNA template are in blue. The RNA primer is in red. The σ subunit domains are shown as rectangles in blue (σ2-3) and green (σ4). The active site registers are indicated as “i” and “i+1”. (B). Scheme of events during transition from ITC to elongation complex (EC). The upper images shows productive ITCs and paused ITCs (PITC) in complex with the LDT scaffold. Bases corresponding to the SDT scaffold are shown in blue. ITCs are shown in the pre-translocated (ITC^Pre^) and post-translocated (ITC^Post^) states. The gray gradients in the lower part of the panel indicate the probability level of abortive transcription and initial pausing, and the accomplishment levels of RNA:DNA hybrid formation, σ displacement and ejection.

### Two steps in the maturation of RNA:DNA hybrids and DNA scrunching

Our results underline two phases in initial transcription: (1) transition from unstable to stable RNA:DNA hybrids when RNA length reaches 5 nt; and (2) transition from ITC prone to pause to productive ITC/EC when RNA length exceeds 8 nt (**Figure 7B**). Before the first phase, abortive transcription is predominant and the RNA:DNA hybrid stability depends on σ. Before the second phase, pausing is predominant and translocation depends on σ. These phases perfectly fit with the three ITC types observed at natural promoter templates: unstable ITCs with RNAs < 4 nt, intermediate stability ITCs with RNAs of 4-8 nt, and stable productive ITC with RNAs > 8 nt (Metzger et al, 1993)(Carpousis & Gralla, 1980). Our results on RNA:DNA hybrid stability are in agreement with structural studies on eukaryotic RNAPII showing that ITCs with 4-5-nt RNAs form distorted or mismatched RNA:DNA hybrids, while the 6-8 bp hybrids harbor identical canonical structures (Liu et al, 2011). Unlike ITCs formed on promoters, ITCs formed on our DNA scaffolds lack the upstream part of the transcription bubble and non-template ssDNA. Therefore, the initial transcription on scaffold is not affected by the stress arising due to DNA scrunching, which is a major cause of abortive transcription (Kapanidis et al, 2006). The lack of stress in DNA templates may explain the quantitative retention of 4-7-nt RNAs in σ-containing ITCs formed on scaffolds, compared with the a small fraction of 5 to 7-nt RNAs retained in ITCs formed on promoter DNA templates (Brodolin et al, 2004)(Duchi et al, 2016).

### σ4 versus σ3.2-finger

We found that the region 4.2 of the σ subunit stimulates *de novo* initiation, stabilizes short RNA:DNA hybrids, and selectively stimulates UTP addition to the RNA 3’ end. These activities were previously attributed to the σ3.2-finger (Kulbachinskiy & Mustaev, 2006)(Pupov et al, 2014)(Zenkin & Severinov, 2004) that stabilizes template ssDNA in the active site (Zhang et al, 2012)(Tupin et al, 2010). Altogether, our results show that σ4.2 activity during initial transcription can be clearly differentiated from that of the σ3.2-finger. Indeed, σ3.2-finger deletion abolished RNA retention in ITCs, but had only a moderate effect on translocation. Conversely, σ4 deletion strongly affected RNA retention and also translocation. Furthermore, the σ3.2-finger was dispensable for *de novo* initiation on scaffold template harboring a long dwDNA duplex, while σ4.2 was essential. Therefore, we conclude that σ4 interaction with core RNAP is a major determinant of the σ subunit transcription-stimulatory activities. Our results also suggest that the σ3.2 and σ4 regions are functionally connected, and changes in σ3.2 structure may affect σ4 function. Therefore, the σ3.2-finger deletion might have an impact on promoter-dependent transcription via allosteric changes in σ4 conformation or/and its positioning in the RNA exit channel. Finally, our results provide a rational explanation to the finding that defects caused by σ3.2-finger deletion are promoter-sequence dependent (Kulbachinskiy & Mustaev, 2006)(Morichaud *et al*, 2016).

### The σ subunit domain 4 as allosteric regulator of NAC

The domain σ4 does not make any contact with the scaffold DNA or RNA, and therefore exerts its anti-pausing activity through interaction with the RNAP core. In the RNAP holoenzyme, the region 4.1 of σ4 is located in the RNA exit channel and occupies the place of RNA, 13-16 nucleotides from the RNA 3’ end. Region 4.2 of σ4 interacts with β-FTH. Therefore, σ4 may influence translocation, like the RNA structures formed within the RNA-exit channel that can affect RNAP active site conformational cycling (trigger loop folding/unfolding) through β-flap interaction (Toulokhonov & Landick, 2003)(Toulokhonov *et al*, 2001) (Hein *et al*, 2014). In addition, σ4.2 interaction with the Zn-binding domain of the β’-clamp might influence clamp conformational motions during translocation. Paused elongation complexes are characterized by several changes in RNAP structure: rotation of the swivel module (comprising the β’ clamp), disordered β-flap (Kang *et al*, 2018), widened RNA exit channel, and partially open clamp state (Weixlbaumer *et al*, 2013). The PITC complexes formed in our assay likely resemble the crystallized elemental paused elongation complexes (ePEC) formed with scaffolds the architecture of which was almost identical to ours (lacking non-template strand ssDNA) and that were trapped in a partially open clamp conformation (Weixlbaumer *et al*, 2013). We hypothesize that in the absence of 9-bp RNA:DNA hybrids and σ4, RNAP may be blocked in the open-clamp/non-swiveled state, thus inhibiting forward translocation. The σ4 interaction with the RNA exit channel/clamp may promote conformational motions of the swivel module/clamp and thus stimulate translocation during initial transcription.

### Analogy in function of the basal transcription factors from different kingdoms of life

We demonstrated that the anti-pausing activity can be observed with structurally distinct σ subunits and phylogenetically distant RNAPs (*E. coli* and *M. tuberculosis*). Considering that the σ4 and β-flap interaction is invariant between all classes of σ subunits, we propose that σ anti-pausing activity is an universal feature of initial transcription in bacteria. We hypothesize that the function mechanism of archaeal TFB and eukaryotic TFIIB, which are implicated in RNA synthesis priming, might be similar to that of the σ subunit. Indeed, TFIIB, in which B-reader and B-ribbon are topological homologues of σ3.2 and σ4 respectively (Liu *et al*, 2010)(Kostrewa *et al*, 2009), can stabilize 5-nt RNA in ITC (Bushnell *et al*, 2004) and stimulate *de novo* transcription initiation on scaffold templates (Sainsbury *et al*, 2013).

## MATERIALS AND METHODS

### Proteins, DNA templates, and RNA oligonucleotides

Recombinant core RNAP (harboring the C-terminal 6xHis-tag on rpoC) of *E. coli* (expression plasmid pVS10) and *M. tuberculosis* (expression plasmid pMR4) were expressed in *E. coli* BL21(DE3) cells and purified as described before (Hu *et al*, 2014) (Morichaud *et al*, 2016). The σ^70^ and σ^B^ subunits and their mutants (harboring an N-terminal 6xHis-tag) were constructed and produced as described (Hu *et al*, 2014) (Morichaud *et al*, 2016)(Zenkin *et al*, 2007). RNA oligonucleotides were purchased from Eurogentec and DNA oligonucleotides from Sigma-Aldrich. All oligonucleotides were HPLC purified. AsiA was a generous gift from Dr. Deborah Hinton. The 116bp *lac*UV5 was prepared by PCR amplification (Morichaud *et al*, 2016). The 72-bp *gal*P1cons promoter fragment (promoter positions −50 to +22) was prepared by annealing two oligonucleotides (upper strand oligonucleotide labeled by Cy3 at the 5’ end: 5’-GTTTATTCCA TGTCACACTT TTCGCATCTT TTCGTTGCTA TAATTATTTC ATACCAAAAG CCTAATGGAG CG-3’, and bottom strand: 5’-CGCTCCATTA GGCTTTTGGT ATGAAATAAT TATAGCAACG AAAAGATGCG AAAAGTGTGA CATGGAATAA AC-3’) followed by purification on 10% PAGE. To assemble DNA scaffolds, oligonucleotides were heated in transcription buffer (TB; 40 mM HEPES pH 7.9, 5mM MgCl_2_ 50 mM NaCl, 5% glycerol) at 65°C for 5 min, and then annealed by lowering the temperature to 16°C for 30 min.

### Scaffold-based transcription assays

Transcription reactions were performed in 5 μl of TB. 240 nM RNAP core was mixed with 1 μM of full length σ or 2μM of σ mutants, and incubated at 37°C for 5 min. 1μM RbpA was added when indicated. Samples were supplemented with 0.8 μM (final concentration) scaffold DNA and 50 μM (final concentration) pRNA. The primer extension kinetics were evaluated using 100 μM pRNA. Samples were incubated on ice for 5 min, then at 22°C for 5 min, and supplemented with 0.4 μM [α-^32^P]-UTP or [α-^32^P]-CTP and 22 μM of the indicated NTPs (HPLC purified). Transcription was performed at 22°C for 2 min or for the indicated time and stopped by adding an amount of loading buffer (8M Urea, 50mM EDTA, 0.05% bromophenol blue) equal to the reaction volume. Samples were heated at 65°C for 2 min and RNA products were resolved on 26% PAGE (acrylamide : bis-acrylamide ratio 10:1) with 7M urea and 1x TBE.

### RNA retention assay

*Eco*RNAP-scaffold DNA-RNA complexes were assembled as described above except that 100 μM of 2-3-nt pRNA and 10 μM of 4-5-nt pRNA were used. pRNA7 and pRNA13 were mixed with scaffold DNA before annealing. Complexes formed in 5μl TB in Axygen® 1.7 ml MaxyClear Microtubes were incubated at 18°C for 5 min. Then, 5μl of Ni-NTA agarose beads slurry (Qiagen) in TB was added, and tubes shaken using an Eppendorf ThermoMixer^®^ at 18°C for 5 min. To separate the Ni-bound RNAP fraction, 0.5 ml of TB/250mM NaCl was added. Samples were briefly stirred, pelleted by centrifugation at 1000g for 1 min, and supernatants were discarded. A second washing step was performed with 50 μl of TB as before. Supernatants were removed to leave a sample volume of 10μl. [α-^32^P]-UTP (0.4 μM final concentration) was added to all samples that were then incubated at 22°C for 3 min. Reactions were quenched and analyzed as above.

### Promoter-based transcription assays

Transcription on the *lac*UV5 and *gal*P1cons promoters was performed with 200 nM *Eco*RNAP, 500 nM full length σ^70^ or 2μM σ^70^ mutants, and 300 nM DNA template in TB. Samples were incubation at 37° for 10 min, and supplemented with 100 μM ApA (*lac*UV5 assay) or CpA (*gal*P1cons assay) and 0.4 μM [α-^32^P]-UTP. Transcription reactions were performed at 37°C for 10 min.

### AsiA inhibition assay

500 nM σ^70^ was first mixed with 1 μM AsiA and then with 100 nM *Eco*RNAP core. Samples were incubated at 30°C for 10 min. Next, DNA templates were added and samples were incubated at 30°C (with *lac*UV5 promoter) and at 22°C (with SDT2 scaffold) for 5 min. Transcription from the *lac*UV5 promoter (50 nM) was initiated by adding 100 μM ApA and 0.4 μM [α-^32^P]-UTP at 37°C for 5 min. Transcription from scaffold DNA (0.8 μM) was performed in the presence of 0.4 μM [α-^32^P]-UTP and 25 μM NTPs or 50 μM GpC and carried out at 22°C for 3 min.

### Calculation of the pause half-life times

The pause half-life times (t_1/2_ = *ln2/k*) were calculated by fitting the fractions of RNA in pause 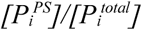 in function of the time *t* using the following single-exponential equation: 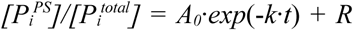, where 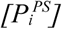 is the RNA in pause at the time point *i* and 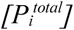 is the total RNA at the time point *i. A* is the amplitude and *R* is the residual.

## ACKNOWLEDGMENTS

We thank Georgiy Belogurov for discussion of the results and Nikolay Zenkin for critical reading. We are grateful to Deborah Hinton for providing AsiA protein. Funding was from the French National Research Agency [MycoMaster ANR-16-CE11-0025-01].

## AUTHOR CONTRIBUTIONS

K.B conceived the study and designed exepriments, K.B. and Z.M. performed experiments. K.B. performed data analysis and wrote the manuscript with contribution from Z.M.

