## Supplemental Figures 1 to 6 for "Anti-pausing activity of region 4 of the RNA polymerase σ subunit and its regulation by σ-remodeling factors"

IRIM, CNRS, Univ Montpellier, 1919 route de Mende 34293 Montpellier, France

**Supplementary figures and legends**

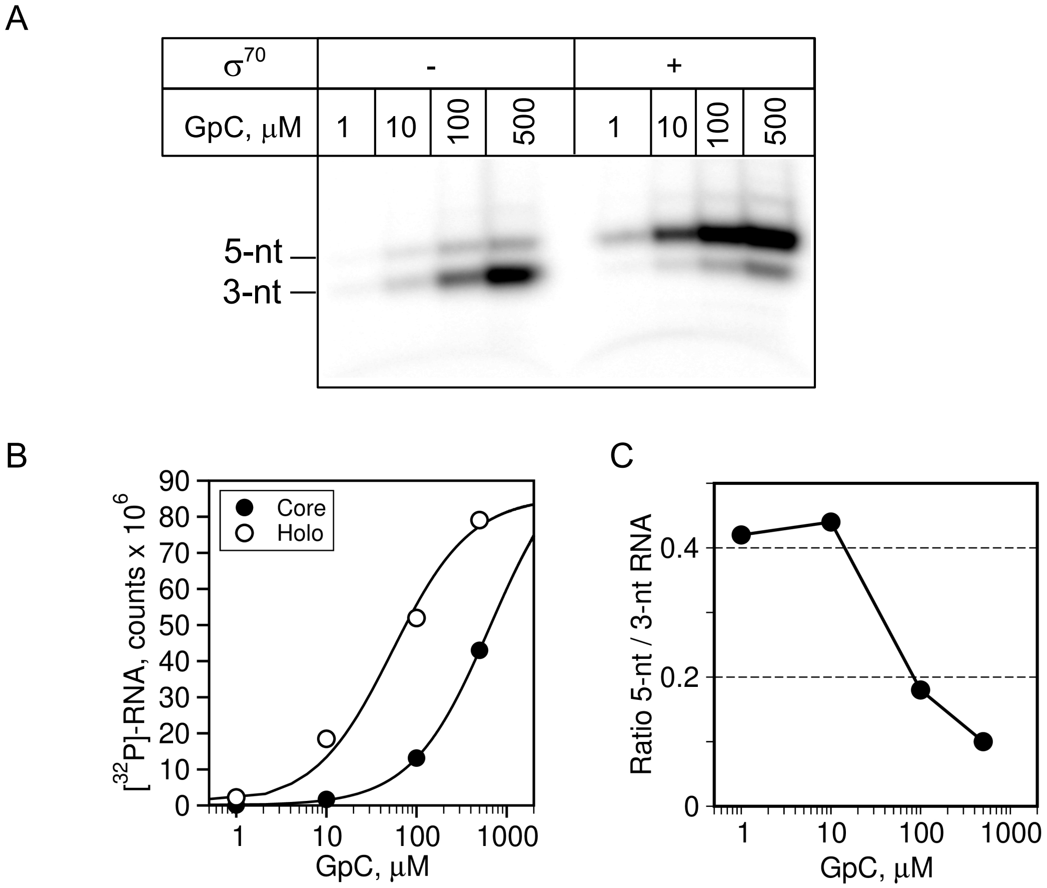


**Figure S1. Effect of the pRNA concentration on translocation efficiency.**
(A). Primer extension by the *Eco*RNAP core and $\sigma^{70}$-*Eco*RNAP holoenzyme performed at the SDT scaffold with indicated concentrations of pRNA (GpC) and fixed concentration of [$\alpha^{32}$P]-UTP, GTP and ATP. Transcription was performed for 2 min. (B) Quantification of the experiment shown in panel A. Cumulative RNA products are plotted as function of pRNA concentration. (C). Ratio of the 5-nt to 3-nt RNAs for reactions with *Eco*RNAP core plotted as function of pRNA concentration.

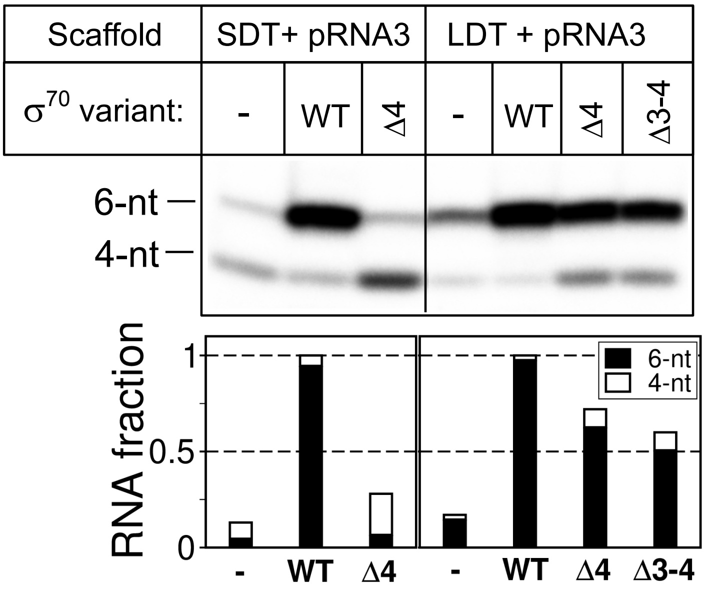


**Figure S2. The dwDNA duplex promotes forward translocation.**

Extension of 3-nt pRNA (pRNA3) by *Eco*RNAP on the SDT and LDT scaffolds in presence of the mutant $\sigma^{70}$ variants. Stacked bar graph shows quantification of the RNA products. Values of each RNA product were normalized to the total [$^{32}$P]-RNA synthesized in the presence of the full length $\sigma^{70}$ subunit.

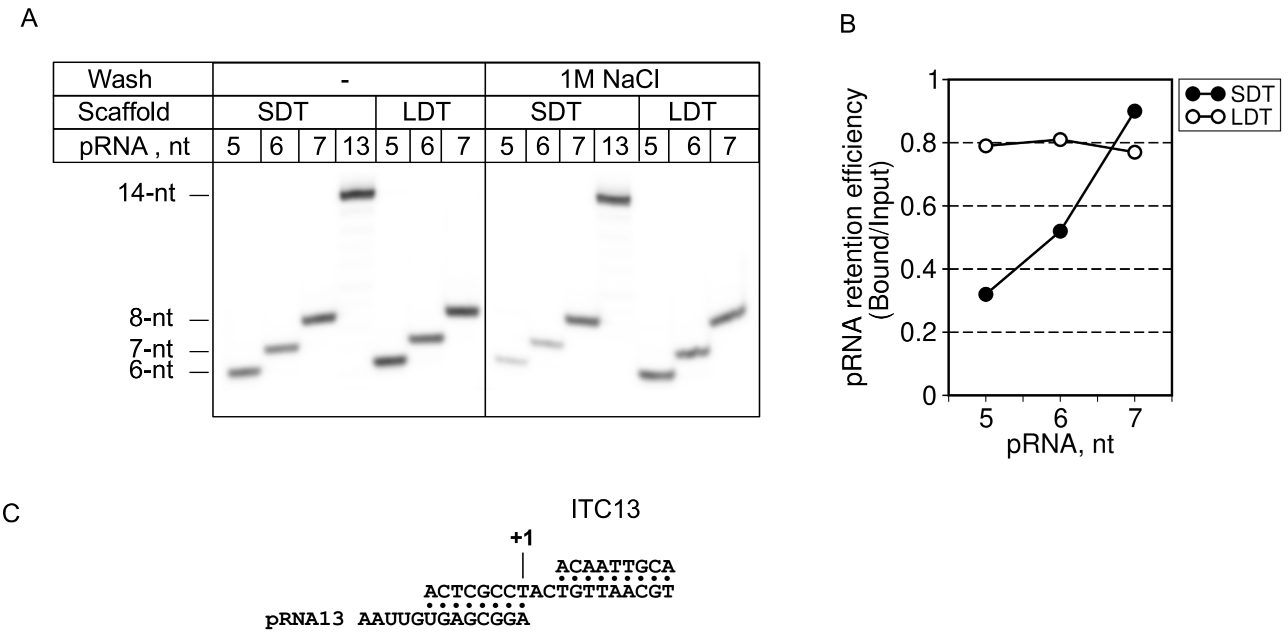


**Figure S3. RNA retention in ITCs formed by the *Eco*RNAP core.**

(A). [$\alpha^{32}$P]-UTP labeled RNAs produced in primer extension reactions in ITC5, ITC6, ITC7 (5- 7-nt pRNA shown in **Fig. S4A**) and ITC13 (shown in panel C) assembled on SDT and LDT scaffolds. ITCs were immobilized on $Ni^{2+}$-agarose beads and washed with buffer containing 1M NaCl. (B). The fraction of labeled RNA retained in ITCs in panel A was calculated as a ratio between RNA in the washed ITCn to the RNA in the unwashed ITCn. (C). Scheme of the SDT scaffold with 13-nt pRNA used to assemble ITC13.

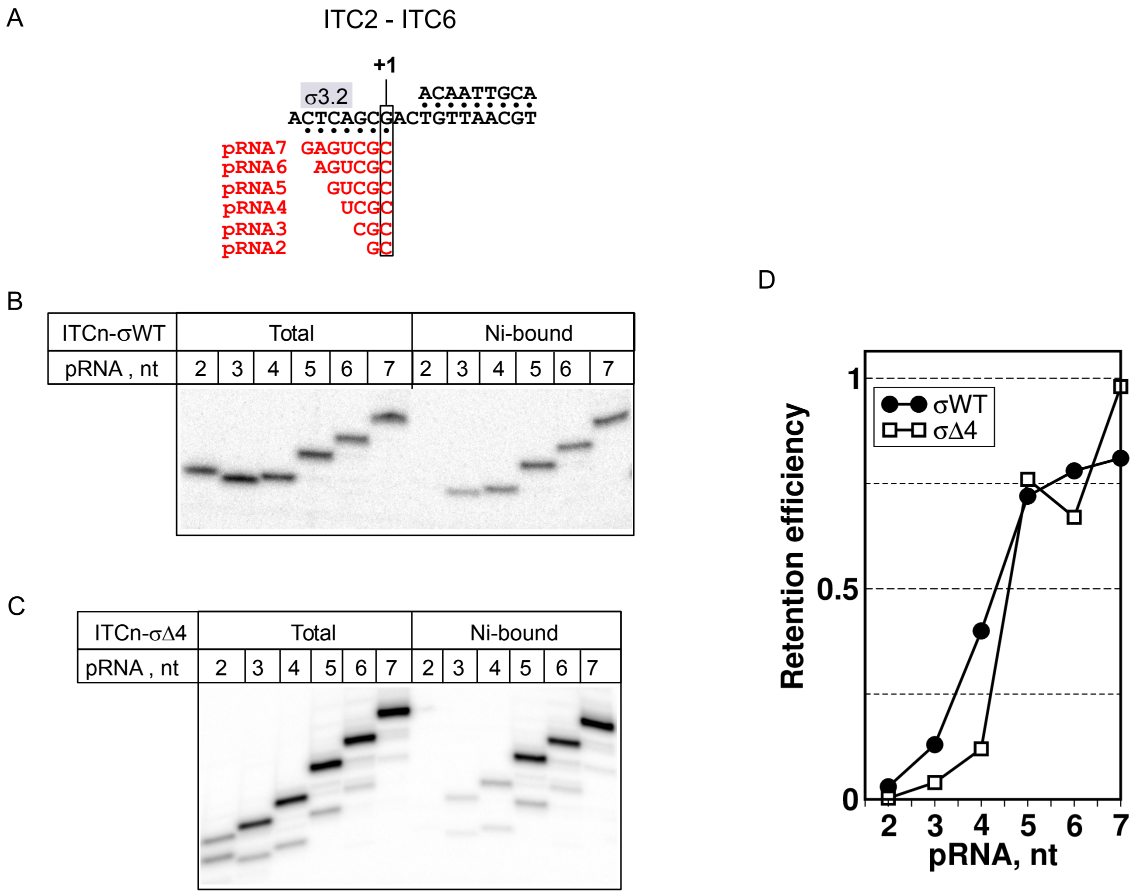


**Figure S4. The σ subunit region 4 stabilizes short RNAs in ITCs formed on SDT scaffold.**

(A). Scheme of the SDT scaffold with RNA primers used to assemble ITC2 - ITC7. (B and C). [$\alpha^{32}$P]-UTP labeled RNAs produced in primer extension reactions in the presence of the $\sigma^{70}$ (EσWT) and σΔ4 (EσΔ4) respectively. ITCs were immobilized on $Ni^{2+}$-agarose beads and washed with buffer, when indicated, before labeling. For the EσΔ4, GTP and ATP were added to the reactions (E). Quantification of the panels B and C. The fraction of labeled RNA retained in ITCn was calculated as a ratio between RNA in the washed ITCn to the RNA in the unwashed ITCn.

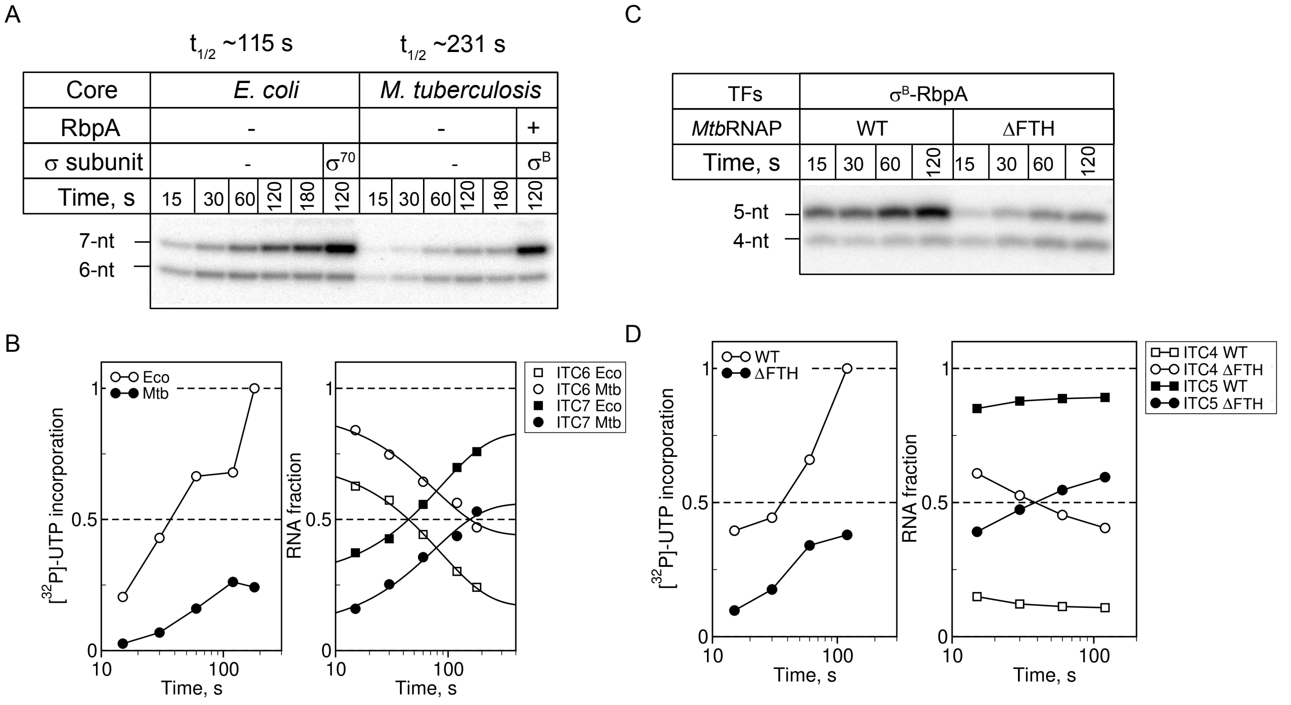


**Figure S5. $\sigma^B$ and β-flap modulate the initial pause.**

(A). Kinetics of the 5-nt pRNA extension by the RNAP core from *E. coli* and *M. tuberculosis* at the SDT2 scaffold. Transcription was initiated by [$\alpha^{32}$P]-UTP and GTP. **(B).** Quantification of the experiment shown in panel A. Left graph shows synthesis of total [$^{32}$P]-RNA as function of time. Right graph shows fractions of [$^{32}$P]-RNA products, representing various ITCs, as function of time. For each time point, values were normalized to the total [$^{32}$P]-RNA synthesized at that time point. (C). Kinetics of the 3-nt pRNA extension by the wild type $\sigma^B$-*Mtb*RNAP (WT) and mutant $\sigma^B$-*Mtb*RNAP$^{\Delta FTH}$ (ΔFTH) holoenzyme at SDT2 scaffold in the presence of RbpA. Transcription was initiated by [$\alpha^{32}$P]-UTP and GTP. (D). Quantification of the experiment shown in panel A the same way as in panel B.

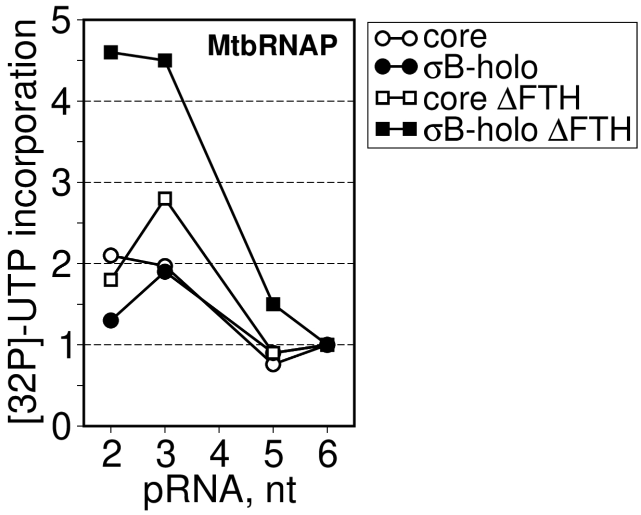


**Figure S6. The $\sigma^B$ subunit increases abortive transcription with the mutant *Mtb*RNAP$^{\Delta FTH}$.**

Quantification of the RNA products shown in Fig. 6F. For each condition, RNA in each lane was normalized to RNA synthesized in ITC7.
